# Stratigraphic Paleobiology of Carbonate Systems

**DOI:** 10.64898/2026.06.12.732006

**Authors:** Niklas Hohmann, Sidney Bickerton, Anna Jansen, Xianyi Liu, Emilia Jarochowska

## Abstract

Stratigraphic paleobiology is a newly established interdisciplinary approach, which has demonstrated that the fossil record is a joint expression of biotic and stratigraphic change, and all inferences from it must be grounded in a solid understanding of the stratigraphic context. Fossiliferous strata can be found in all depositional systems (e.g., marine, terrestrial, or lacustrine; siliciclastics or carbonates), each having a unique characteristic timescale and set of external controls, which govern the accumulation of sedimentary particles, including fossils. Consequently, the same biotic changes are preserved differently across depositional systems.

While carbonate systems form a large portion of the fossil record, most studies in stratigraphic paleobiology have focused on siliciclastic systems and are not easily generalizable. As they are predominantly formed by living organisms, carbonates are both fossils and record, opening the opportunity to study the co-dependency of life and its environment.

Here, we explore the stratigraphic paleobiology of carbonate systems by combining simulations of carbonate platform and ramp geometries with synthetic fossil records. We explore the preservation of extinction patterns and rates spatially and across geometries. By examining stratigraphic biases in isolation (unconformity and condensation, ecology, and abundance biases), we find characteristic differences between ramp and platform geometries due to their differential response to sea level change, spatial variability, and differences in ecological clines.

Differences in the structure of the fossil record between platform geometries are traceable to the contribution and properties of the carbonate producing organisms (carbonate factories), showing that preservation of earth system data in carbonate systems will vary both latitudinally and temporally or as a result of major perturbations of the biogeosphere. Our results show that while general rules on the structure of the fossil record can be derived for entire depositional systems, accounting for the geological and ecological dynamics of a particular sedimentary basin can hugely refine interpretations of the fossil record. That is particularly true for biogenic and biologically-mediated sediments.

## Introduction

Stratigraphic paleobiology posits that the fossil record is a joint expression of sedimentological, stratigraphic, and biotic change, and that any reading of the fossil record must be grounded in a solid understanding of local stratigraphic context and the response of stratigraphic architectures to their external controls. The importance of the stratigraphic context for a plethora of paleodisciplines, including, but not limited to, paleoecology, macroevolution, and phenotypic evolution, has been demonstrated by many studies (Holland and Patzkowsky, 1999; Hannisdal, 2006; Peters, 2008; Danise *et al*., 2019; Zimmt *et al*., 2021; Hohmann *et al*., 2024). A prime example of this is the insight that four of the “big five” mass extinctions are associated with major fluctuations in sea level, which artificially inflates extinction rates due to extirpation of niche-tracking taxa due to water depth changes across unconformities and condensation (Holland and Patzkowsky, 2015; Holland, 2020).

Understanding the stratigraphic distribution of fossils is at the core of the stratigraphic paleobiology research program. Both in empirical and in simulation studies, a large portion of stratigraphic paleobiology focuses on the identification of artefactual peaks in origination and extinction rates due to unconformities and extirpation (Holland and Patzkowsky 2015; Nawrot et al. 2018), as well as on paleoecology and biotic interactions (Scarponi and Kowalewski, 2004; Huntley and Scarponi, 2015; Mondanaro, Dominici and Danise, 2024).

More generally, fossils record a wide range of data on earth and life systems, such as climate, evolution, and geochemistry. Naturally, a stratigraphically biased distribution of fossils will have downstream effects on those study systems, and there has been significant effort to propagate principles of stratigraphic paleobiology into other disciplines drawing upon the fossil record (Holland, Patzkowsky and Loughney, 2025). For example, this includes the study of gaps and transport on isotope records (Curtis *et al*., 2025; van Wieren, Dyer and Husson, 2026) or the recoverability of tempo and mode of phenotypic evolution (Hannisdal, 2006; Hohmann *et al*., 2024), with direct implications for the gradualism vs. punctuated equilibrium debate in evolutionary biology (Eldredge and Gould, 1972; Hunt, Voje and Liow, 2025; Polly, 2025). In other disciplines, such as phylogenetics, uptake remains slow even though first principles have been demonstrated and both simulation and inference tools are available (Heath, Huelsenbeck and Stadler, 2014; Holland, 2016; Barido-Sottani *et al*., 2019; Hohmann and Jarochowska, 2025).

Holland (2000) provides a conceptual partition of the stratigraphic biases jointly structuring the fossil record:

- Unconformity bias, where time intervals are not preserved due to erosion or nondeposition, resulting in data loss (Kabanov, 2017),
- Condensation bias, where unrecognized variations in sedimentation rate lead to the inflation or deflation of rates (Hohmann, 2021),
- Ecological bias (facies bias sensu Holland (2000)), where taxa track their niche, leading to varying fossil abundance and potentially extirpation as clines vary stratigraphically,
- Abundance bias, where low fossil sampling rates can lead to backwards and forwards smearing of extinction and origination rates (Signor-Lipps effect (Signor *et al*., 1982; Foote, 2001)) or mistaking random fluctuations for meaningful observations.

The relative contribution of these biases in structuring the fossil record will inevitably vary across space, time, and between depositional systems.

A majority of early and seminal works in stratigraphic paleobiology addressed marine siliciclastic systems (Holland, 1995, 2000; Nawrot *et al*., 2018; Zimmt *et al*., 2021), with few empirical studies using data from carbonate systems (Holland and Patzkowsky 2009). Recently there has been a push to generalize these results to nonmarine environments (Holland and Loughney, 2021; Loughney *et al*., 2021; Loughney and Holland, 2026) and tectonostratigraphy and macrostratigraphy, covering longer (> 10 Myr) time scales (Holland 2016; Holland et al. 2025a). The unifying paradigm across the discipline is that stratigraphic architectures can be understood in a sequence stratigraphic framework, and the structure of the fossil record can be explained on a systems tracts basis as a function of external drivers, such as subsidence and eustatic sea level that generate accommodation space and ecological clines.

However, it is well documented that the sequence stratigraphy of (marine) temperate and tropical carbonate depositional systems differs significantly from siliciclastic systems (Schlager, 2005). This can be attributed to rapid rates of cementation and high rates of in situ biogenic sediment production, making carbonate systems literally both fossil and record. Depending on the dominant carbonate factories, carbonate systems display a wide array of geometries, ranging from rimmed platforms to ramps (Burchette and Wright, 1992). While categorization of carbonate systems by geometry is a priori purely descriptive, it is intimately connected to their tectonic setting, as well as to the sediment producing and binding organisms that comprise them (Kenter, 1990; Pomar, 2001; Bosence, 2005). As these organisms react differently to changing environmental conditions such as water depth, temperature, and light availability, different carbonate geometries will be displayed based on different latitudinal distributions and responses to external drivers and internal changes (Tucker, Calvet and Hunt, 1993; Pomar, 2001; Lehrmann *et al*., 2022). For example, rapidly growing euphotic, framework-building organisms will quickly fill available accommodation space when light is available, resulting in the formation of platform geometries in the tropics with a clear separation into platform top and distal slope deposits. In contrast, mesophotic organisms producing coarse sediment will produce ramp geometries much more resembling siliciclastic geometries when carbonate saturation is low and cementation, accordingly, slow (Pomar, 2001; Adams and Kenter, 2014; Lehrmann *et al*., 2022). This dynamic interaction between biotic carbonate producers and the structure of the sedimentary record can give rise to interesting, yet underexplored feedback loops between life and its record, e.g., where large scale perturbations of the biogeosphere fundamentally change the nature of the fossil record. For example, the frequency of ramp geometries peaks after major mass extinctions (Burchette and Wright, 1992), and Kammer and Ausich (2006) attribute a spike in crinoid biodiversity in the Mississippian to increased circulation due to the collapse of platform-edge reefs during the Late Devonian mass extinctions and the resulting development of platform geometries.

Carbonate rocks form a large portion of the fossil record. For example, Balseiro and Powell (2023) found that in North America, carbonates are twice as densely sampled as siliciclastics, suspecting a preference to sample them due to their increased diversity and the resulting ease of describing new fossil taxa. The volume of preserved carbonate rocks drops after the Paleozoic, potentially being a determinant of major changes in the composition of marine biota throughout the Phanerozoic (Sepkoski, 1981; Peters, 2008).

Combined, understanding the idiosyncrasies of carbonate depositional systems and their stratigraphic paleobiology is crucial for a stratigraphically robust interpretation of a large portion of the fossil record.

Here, we explore the stratigraphic paleobiology of carbonate depositional systems with a focus on the differences between ramp and platform geometries using a novel, bespoke model for carbonate stratigraphic forward modeling (Hidding *et al*., 2025). In addition to examining the preservation of extinction pulses, we partition stratigraphic bias into three subcomponents – unconformity and condensation bias, ecological bias, and fossil abundance bias (Signor-Lipps effect) – to arrive at general principles that apply to all downstream analyses of fossil data from carbonate platforms. We find that the biases from platform geometries are dominated by unconformity and condensation bias with a spatially clear separation into platform top and ramp, while ramp geometries show more pronounced progradational/retrogradational patterns and a more gradual change of unconformity and condensation bias and ecological clines along the onshore/offshore gradient. The differences between these carbonate geometries can be explained by differences in sediment transport, cementation time, and growth rates of carbonate producing organisms. Our results show the co-dependency of life and its record by demonstrating that stratigraphic architectures of carbonates differ significantly under identical external forcing, and will thus yield different records of identical biotic events.

## Methods

We simulated two attached carbonate platforms, each consisting of three facies reflecting different carbonate factories, extracted stratigraphic architectures and environmental clines from them, and compared differential preservation of biotic signals between them and across the onshore-offshore gradient.

### Carbonate forward modeling

We simulated two attached carbonate platforms using the Julia package CarboKitten.jl (v0.6.0) (Bezanson *et al*., 2017; Hidding *et al*., 2025). The first is a tropical carbonate platform dominated by the euphotic (photozoan) carbonate factory, rapid cementation and little transport, the second a temperate carbonate ramp dominated by the oligophotic (heterozoan) carbonate factory, slow lithification and increased transport (Figure 1). Based on their geometry, we refer to the two cases as platform and ramp, respectively, though they are both instances of carbonate platforms. Carbonate production in both simulations is determined by three carbonate factories with identical production profiles (extinction coefficient and saturation intensity), but different maximum production rates and transport coefficients (Table 1, Table 2).

**Figure 1:**
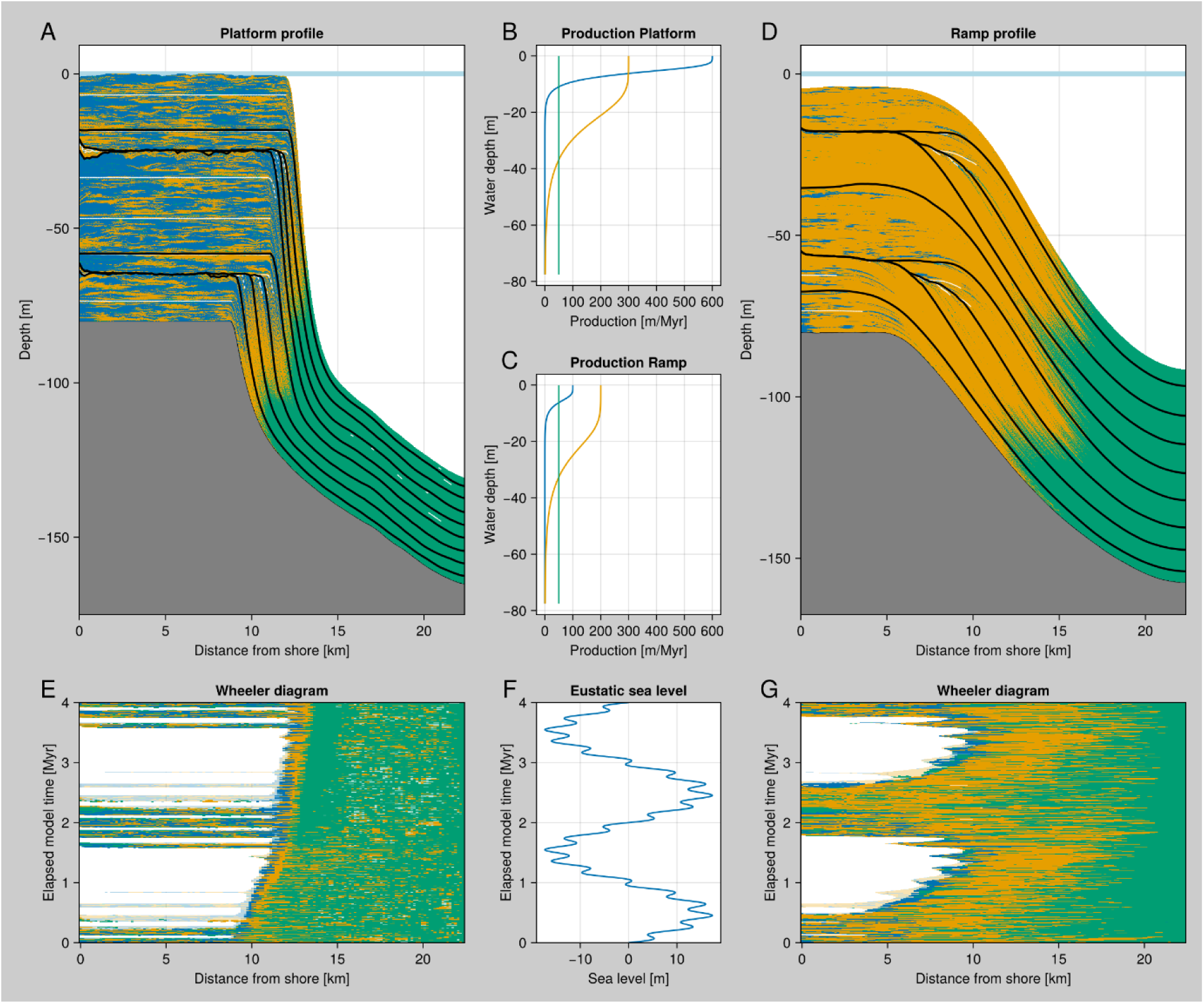
The simulated platform (left) and ramp (right) geometries with profile views (A, D), production curves (B, C), Wheeler diagrams (E, G), and sea level curve (F). See Supplementary Figures 3 and 4 for detailed plots with sampling location and systems tracts.

**Table 1:**
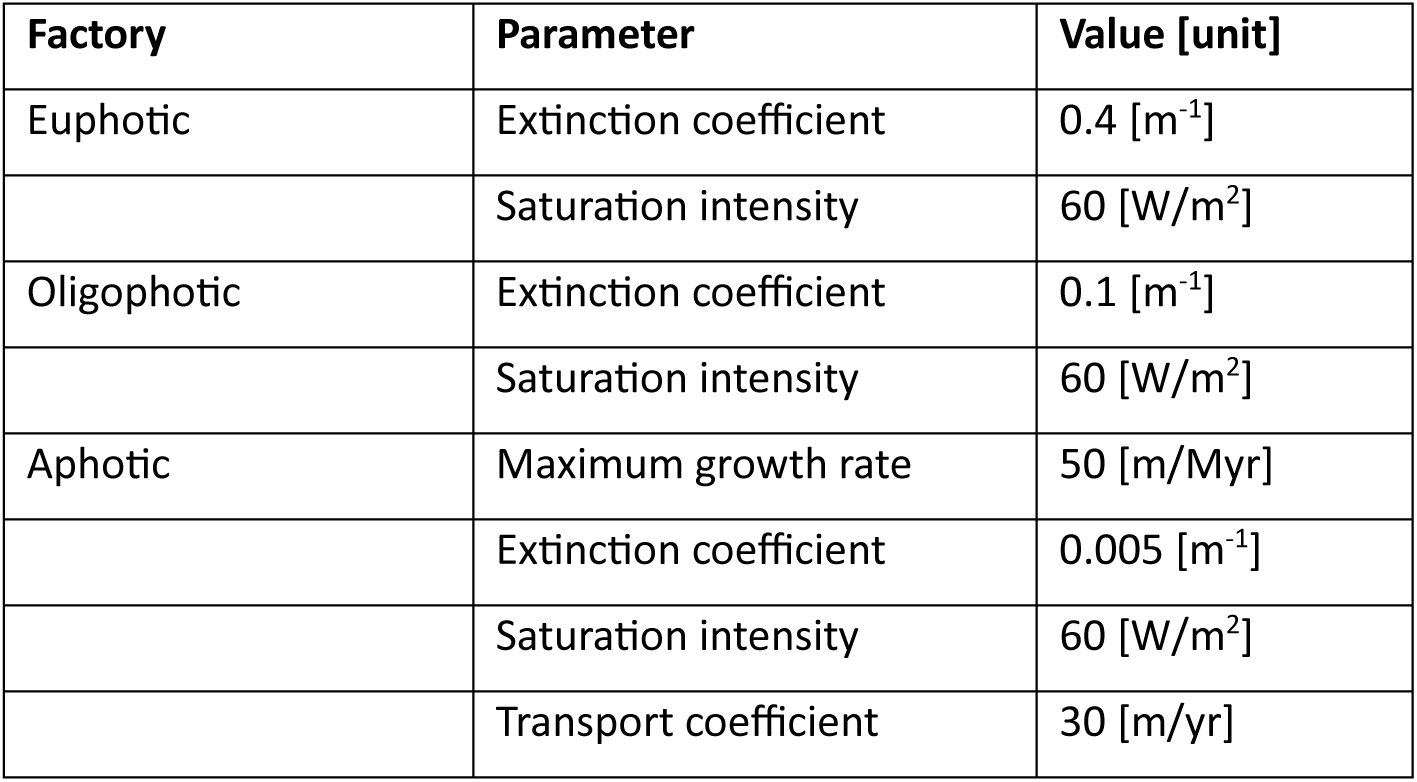
Carbonate factory parameters shared between the platform and ramp simulation. The production-depth curves follow the parametrization by Bosscher and Schlager (1992) (Figure 1).

| Factory | Parameter | Value [unit] |
| --- | --- | --- |
| Euphotic | Extinction coefficient | 0.4 [m<sup>-1</sup>] |
|  | Saturation intensity | 60 [W/m<sup>2</sup>] |
| Oligophotic | Extinction coefficient | 0.1 [m<sup>-1</sup>] |
|  | Saturation intensity | 60 [W/m<sup>2</sup>] |
| Aphotic | Maximum growth rate | 50 [m/Myr] |
|  | Extinction coefficient | 0.005 [m<sup>-1</sup>] |
|  | Saturation intensity | 60 [W/m<sup>2</sup>] |
|  | Transport coefficient | 30 [m/yr] |

**Table 2:** Model parameters differing between the platform geometries.

| Parameter name | Platform | Ramp |
| --- | --- | --- |
| Euphotic maximum growth rate | 600 m/Myr | 100 m/Myr |
| Euphotic transport coefficient | 25 m/yr | 60 m/yr |
| Oligophotic maximum growth rate | 300 m/Myr | 200 m/Myr |
| Oligophotic transport coefficient | 20 m/yr | 60 m/yr |
| Lithification half-life | 0.1 kyr | 2 kyr |
| Precipitation | 2000 mm/yr | 500 mm/yr |

CarboKitten uses a re-implementation of the cellular automaton model by Burgess (2013) to simulate spatial heterogeneity of carbonate deposition due to ecological facilitation, dispersal, and competition of organisms, leading to more empirically realistic spatial and stratigraphic complexity of carbonate strata (Drummond and Dugan, 1999; Burgess and Emery, 2004). Per model time step, a carbonate factory persists in a grid cell when – within a 5 by 5 grid cell neighborhood – the number of grid cells is within viability range, and empty cells are colonized by a factory when the number of neighboring cells is within activation range. We used the default settings for the cellular automaton with an activation range of 6 to 10 and a viability range of 4 to 10 (see Hidding et al. (2025) for more details and discussion).

Variations in carbonate saturation and the resulting differences in time to lithification were modeled via the lithification half-life parameter, which specifies how rapidly the entrained material settles.

Lithification time in the platform was more than an order of magnitude shorter than in the ramp to emulate rapid cementation in tropical carbonate platforms (Table 2).

CarboKitten implements different denudation models. Here, we used the empirical denudation model, assuming precipitation of 2000 mm for the platform and 500 mm for the ramp. This denudation model calculates denudation from local topography and precipitation using an empirical relationship based on multivariate sigmoidal regression of slope and precipitation against denudation calculated from δ^27^Al isotopes (Yang *et al*., 2020).

The subsidence rate in both simulations was set to 20 m/Myr, roughly corresponding to the rates observable in the Bahamas over the Pleistocene (Hearty and Backstrom, 2021), making them moderately aggrading (McNeill, 2005). For the eustatic sea level curve, we used a combination of 3^rd^ and 5^th^ order changes, with periods of 2 and 0.2 Myr and amplitudes of 15 and 2.5 m, respectively. The simulation box was 7.5 km (strike, parallel to shore) by 22.5 km (dip, perpendicular to shore), subdivided into grid cells of 150 m. To reduce artefacts introduced by stratigraphically implausible starting topographies, we used initial topographies generated from separate model runs (pre-runs) over 1 Myr duration under constant sea level (see Supplementary Materials, Supplementary Figures 1 and 2). Main model runs were 4 Myr at a temporal resolution of 100 years, storing data every 1000 years (model resolution).

**Figure 2:**
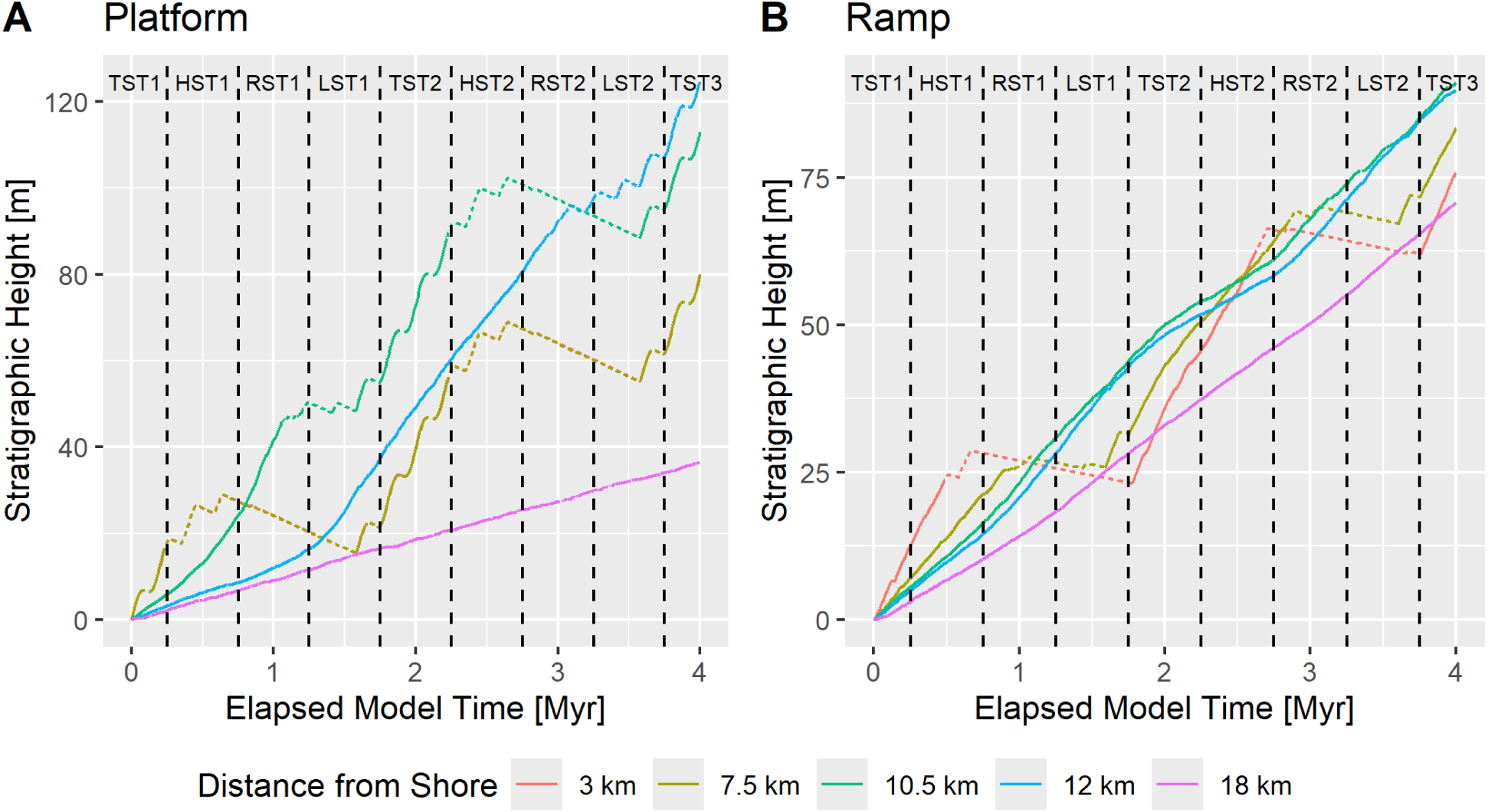
Age-depth models of the platform (left) and ramp (right) geometry. Dashed lines indicate deposition of material later removed and denudation.

Conceptually, we subdivided the simulation time into systems tracts based on the eustatic sea level curve: Transgressive systems tract (TST), highstand systems tract (HST), regressive systems tract (RST), and lowstand systems tract (LST). Each systems tract lasted 0.5 Myr (a quarter of the period of the 3^rd^ order sea level change) and appeared twice during the simulation interval. The exception is the transgressive system tract, which appeared in the middle of the simulation, at its beginning and at its end, but only for 0.25 Myr (Supplementary Figure 3, Supplementary Figure 4). Note that this system-wide sequence stratigraphic subdivision is based on perfect knowledge of the sea level curve, and might differ from the field interpretations made by a sequence stratigrapher based on stacking patterns.

We extracted age-depth models, sediment accumulation curves, and water depth from along-dip transects through the middle of the simulated box and calculated sedimentation rates, as well as summary statistics on hiatus duration (Figure 2, Figure 3, Supplementary Figures 5, 6, 7, 8, 9, 10, and 11). Locations at 3, 7.5, 10.5, 12, and 18 km from the boundary of the simulation box along dip were examined in detail, as they correspond to characteristic locations within their respective depositional geometries (Table 3).

**Figure 3:**
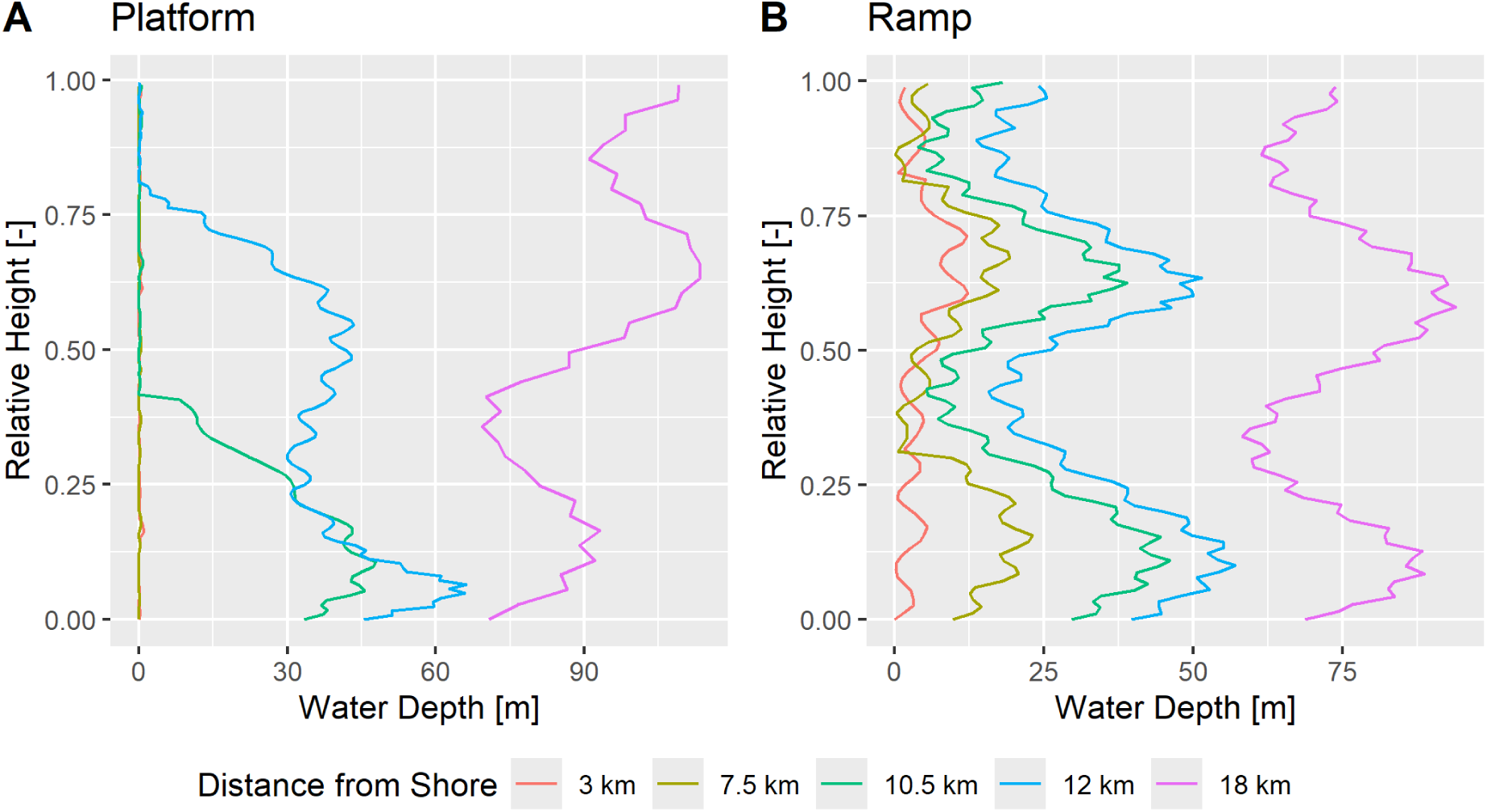
Water depth in the stratigraphic domain. To compare sections of different heights, they were rescaled from 0 (base of section) to 1 (top of section). See Supplementary Figure 12 for water depth in the time domain.

**Table 3:**
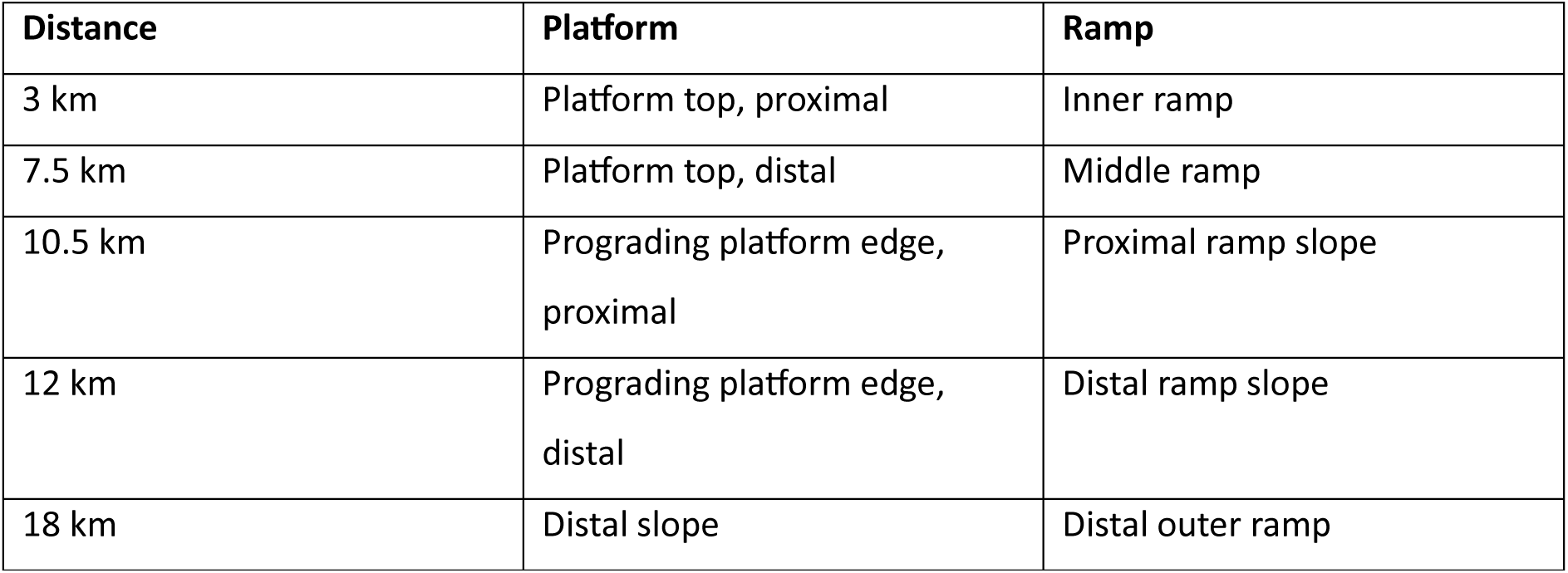
Examined locations in the carbonate platform geometries and their interpretation.

| Distance | Platform | Ramp |
| --- | --- | --- |
| 3 km | Platform top, proximal | Inner ramp |
| 7.5 km | Platform top, distal | Middle ramp |
| 10.5 km | Prograding platform edge, proximal | Proximal ramp slope |
| 12 km | Prograding platform edge, distal | Distal ramp slope |
| 18 km | Distal slope | Distal outer ramp |

Data was further processed in R version 4.5.3 (R Core Team, 2026) using the R packages admtools (Hohmann, 2025; Hohmann *et al*., 2025) for handling of age-depth models and transformations between the stratigraphic and time domain and StratPal (Hohmann, 2024; Hohmann and Jarochowska, 2025) for linking stratigraphy with paleobiology and niche models.

### Ecological bias

We assume water depth is the dominant faunal gradient, but see the discussion for a critical review of this assumption specifically for carbonate systems. We report water depth in the time and stratigraphic domain (Figure 3, Supplementary Figure 12), as downstream effects of water depth will necessarily depend on a taxon’s ecological preferences (preferred water depth and tolerance to fluctuations).

### Unconformity and condensation bias

We grouped unconformity and condensation bias together as age misassignment bias, as their effects are both rooted in the misassignment of fossil ages. Intuitively, age misassignment bias is the misinterpretation of earth systems data due to imperfect knowledge of the underlying true age-depth relationship. To quantify age misassignment bias given a true and an assumed age-depth model, we define the dimensionless condensation ratio (Eq. 1).

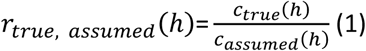

Here, *c_i_*(ℎ) is the condensation at stratigraphic height ℎ, meaning the amount of time recorded per stratigraphic increment under the age-depth model *i* (Hohmann *et al*., 2025). A condensation ratio of 10 over some stratigraphic interval means that, in truth, this interval records 10 times more geological history than inferred based on the assumed age-depth model. This implies that all rates (e.g., origination, extinction, sampling rates, climate change, proxy records – study system independence) calculated on the assumed age-depth model are inflated by a factor of 10. Compared to other measures of age misassignment such as range offset (Holland and Patzkowsky 1999, see below), condensation ratio isolates unconformity and condensation effects, as it only depends on age-depth models. In addition, it directly quantifies how these effects propagate into rates inferred from the stratigraphic record. For this study, we calculated the condensation ratio for the assumed age-depth model of constant, uninterrupted sedimentation, representing a “naive interpretation” of the section where the total duration and thickness of the section is known, but no further stratigraphic or sedimentological information is incorporated (Figure 4).

**Figure 4:**
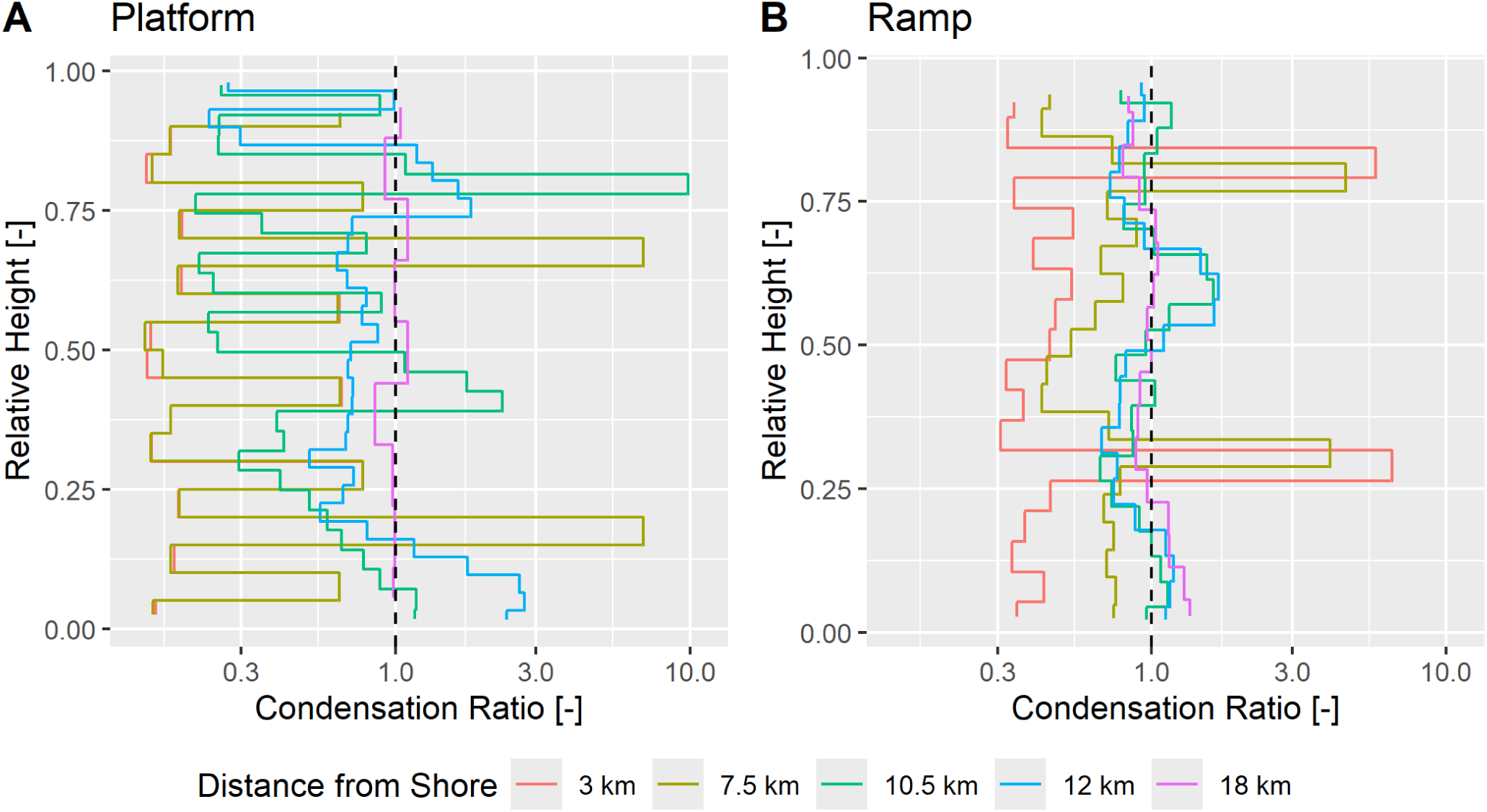
Condensation ratio in the different geometries, measured over stratigraphic intervals of 4 m. To compare sections of differing height, section heights were rescaled from 0 (base of section) to 1 (top of section). The dashed line indicates a condensation ratio of 1, meaning rates are not biased when naïve age-depth relationships are assumed.

### Abundance bias and preservation of extinction pulses

We examined the preservation of pulses of elevated extinction rate in dependence of (1) abundance (sampling rate) of the involved taxa, (2) the timing of the pulse within systems tracts, (3) location along the onshore-offshore gradient and (4) platform geometry (Figure 5).

**Figure 5:**
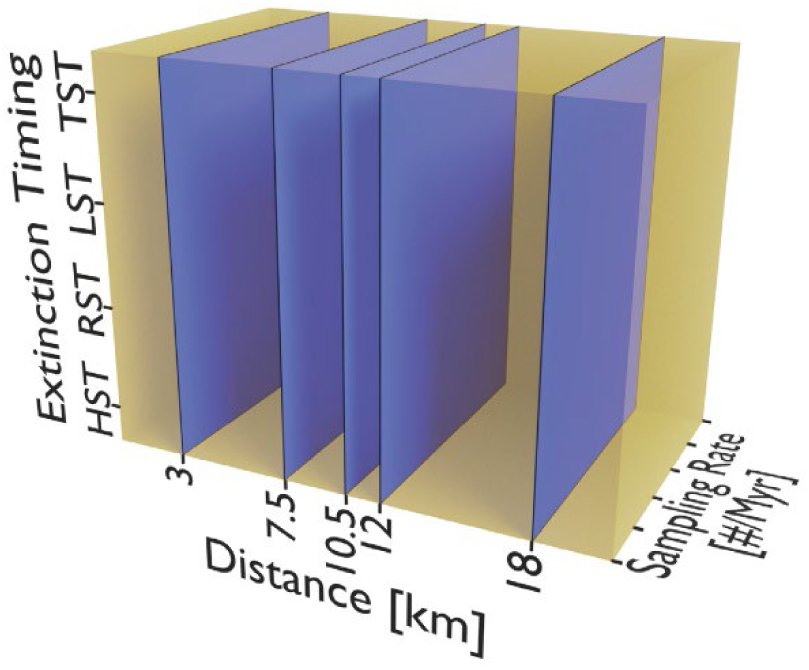
Parameter space for the preservation of extinction pulses within one of the simulated platform geometries. We explore different slices through this three-dimensional space along the depth axis (variations in sampling rate, Figure 6), height axis (position of extinction pulse, Figure 7) and width axis (position along the onshore-offshore gradient, Figure 8 and Figure 9).

We defined five extinction scenarios: One baseline with constant extinction rate, and four scenarios where extinction rate is elevated 25-fold over a constant background rate during the time corresponding to the deposition of one of the systems tracts (HST, TST, LST, and RST, respectively) to mirror the setup by Holland and Patzkowsky (2015) (Supplementary Figure 13). To capture general effects and reduce the influence of stochastic artefacts introduced by small sample sizes, we conditioned simulations on 1000 taxa going extinct. As a result, only relative changes in the rate of last occurrences within a section, but not the rate’s absolute value, is meaningful. The timing of extinction was modeled as a time-variable Poisson point process using the function p3_var_rate from the StratPal package, thus assuming extinctions are independent of each other (Hohmann, 2021). For each taxon going extinct, fossil occurrences up to the time of extinction were simulated using a constant rate Poisson point process with a specified sampling rate using the function p3 of the StratPal package, thus assuming a constant fossil sampling rate in the time domain. We focus on sampling rates ranging from 2 to 100 fossils recovered per Myr. Then, fossil occurrences were transformed into the stratigraphic domain using the age-depth model in which fossils coinciding with gaps (dashed lines in Figure 2) are destroyed. Last occurrences were then determined as the highest preserved fossil per taxon in the stratigraphic domain (Figure 6, Figure 7, Figure 8, Figure 9 and Supplementary Figures 14 to 19).

**Figure 6:**
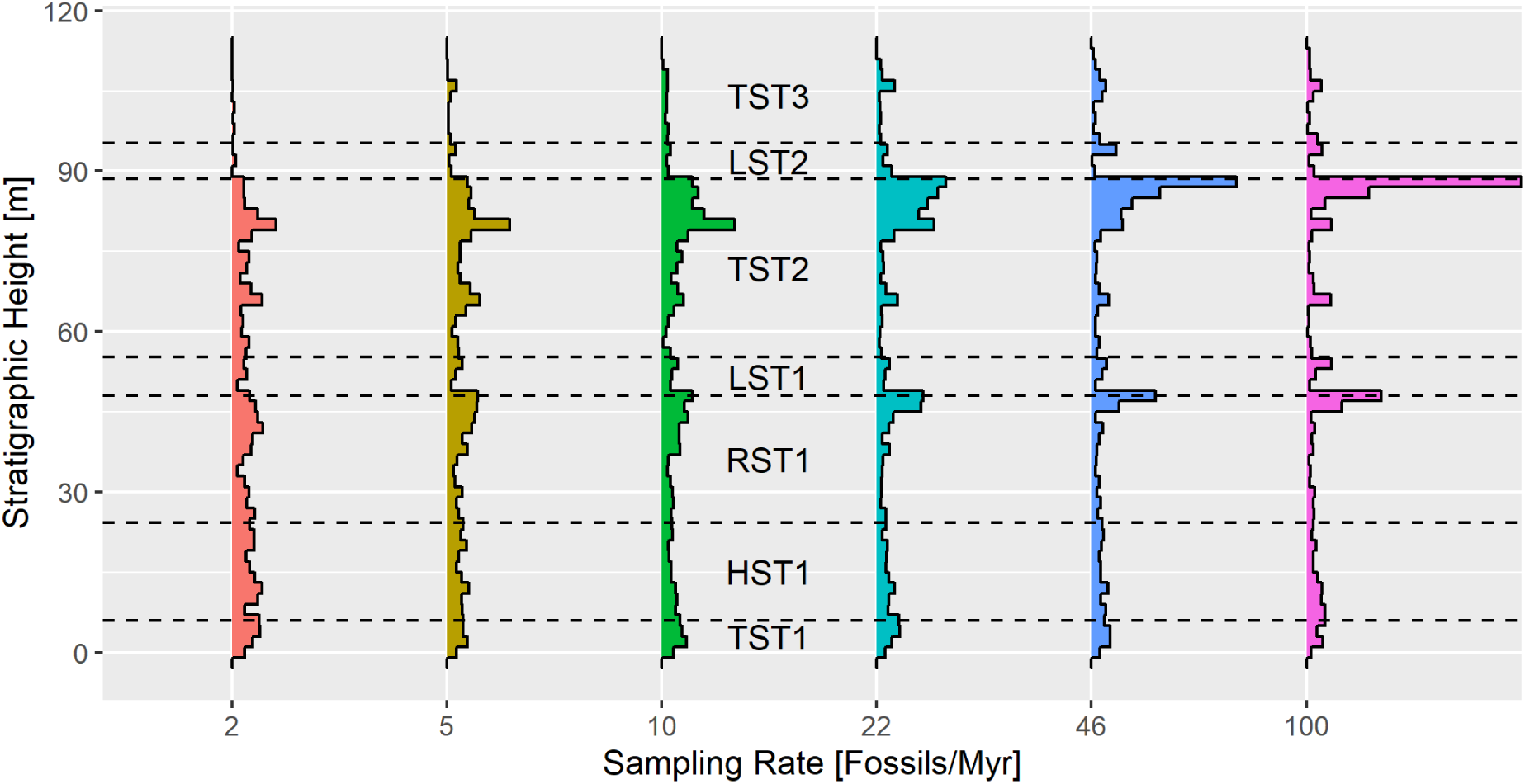
Distribution of last occurrences as a function of fossil sampling rate at 10.5 km from shore (proximal prograding platform edge) on the platform geometry under constant background extinction rate. Results for all other locations are given in Supplementary Figures 14 and 15.

**Figure 7:**
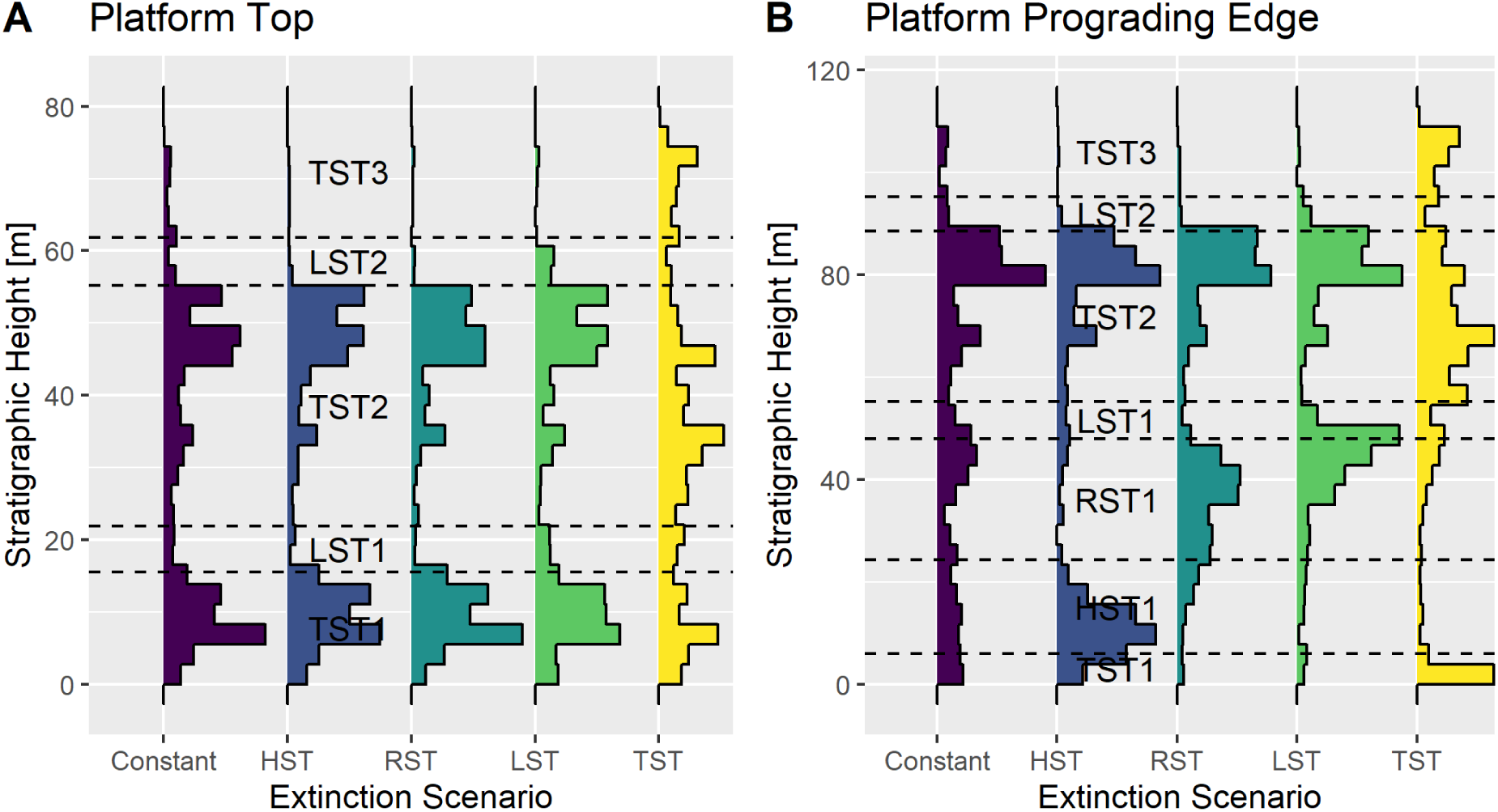
Extinction pulses in different systems tracts and a constant background extinction as baseline preserved (A) on the platform top (3 km from shore) and (B) the proximal prograding edge (10.5 km from shore) in the platform. Fossil sampling rate is 10 fossils per Myr. See Supplementary Figures 16 and 17 for other locations in the ramp and platform.

**Figure 8:**
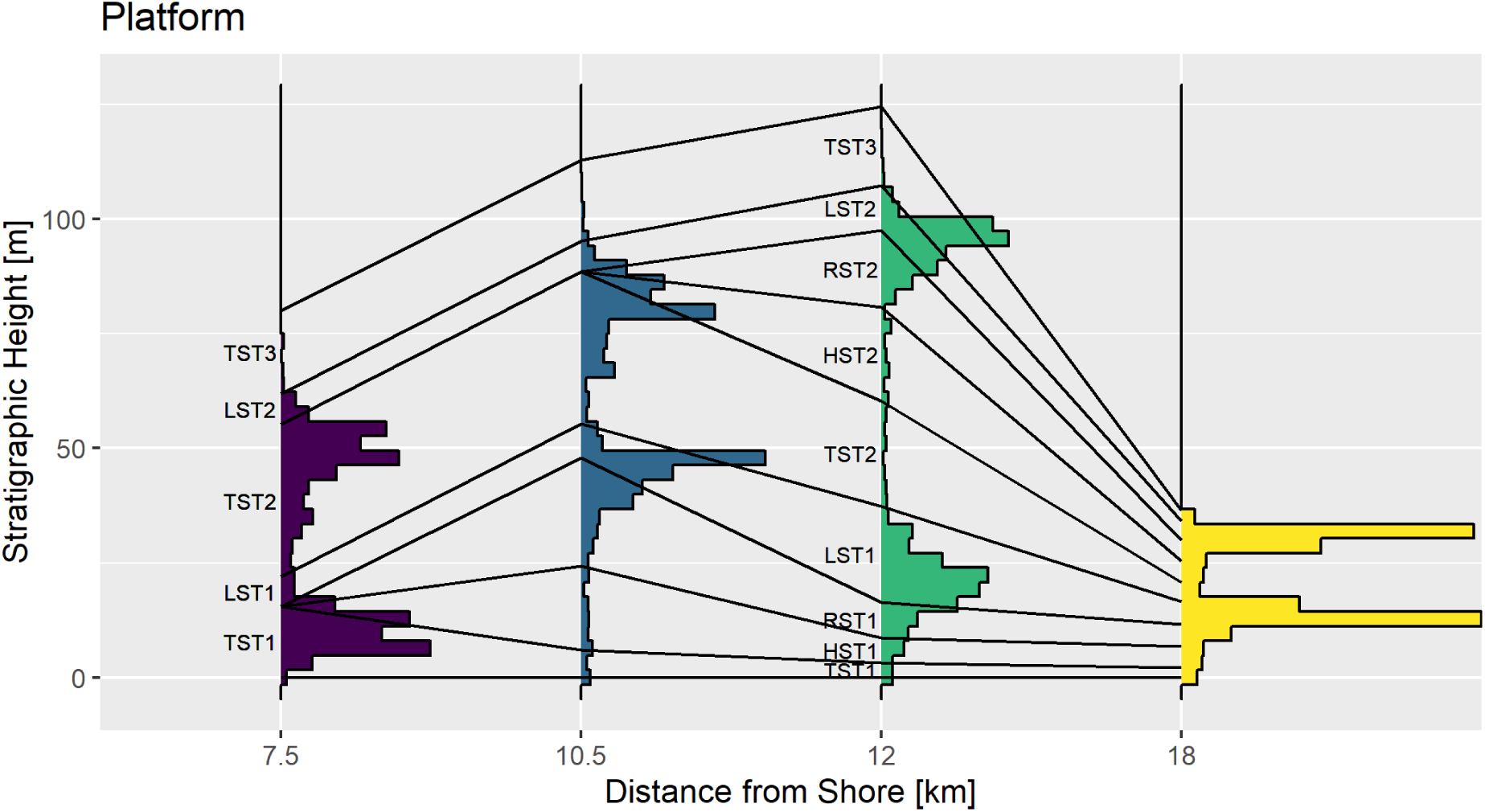
Spatial preservation of a pulse of elevated extinction rate in the lowstand systems tract (LST) in the platform geometry. The proximal platform top at 3 km was omitted as it is identical to the distal platform top at 7.5 km, fossil sampling rate is 10 fossils per Myr. See also Supplementary Figures 18 and 19 for spatial correlation of other extinction pulses.

**Figure 9:**
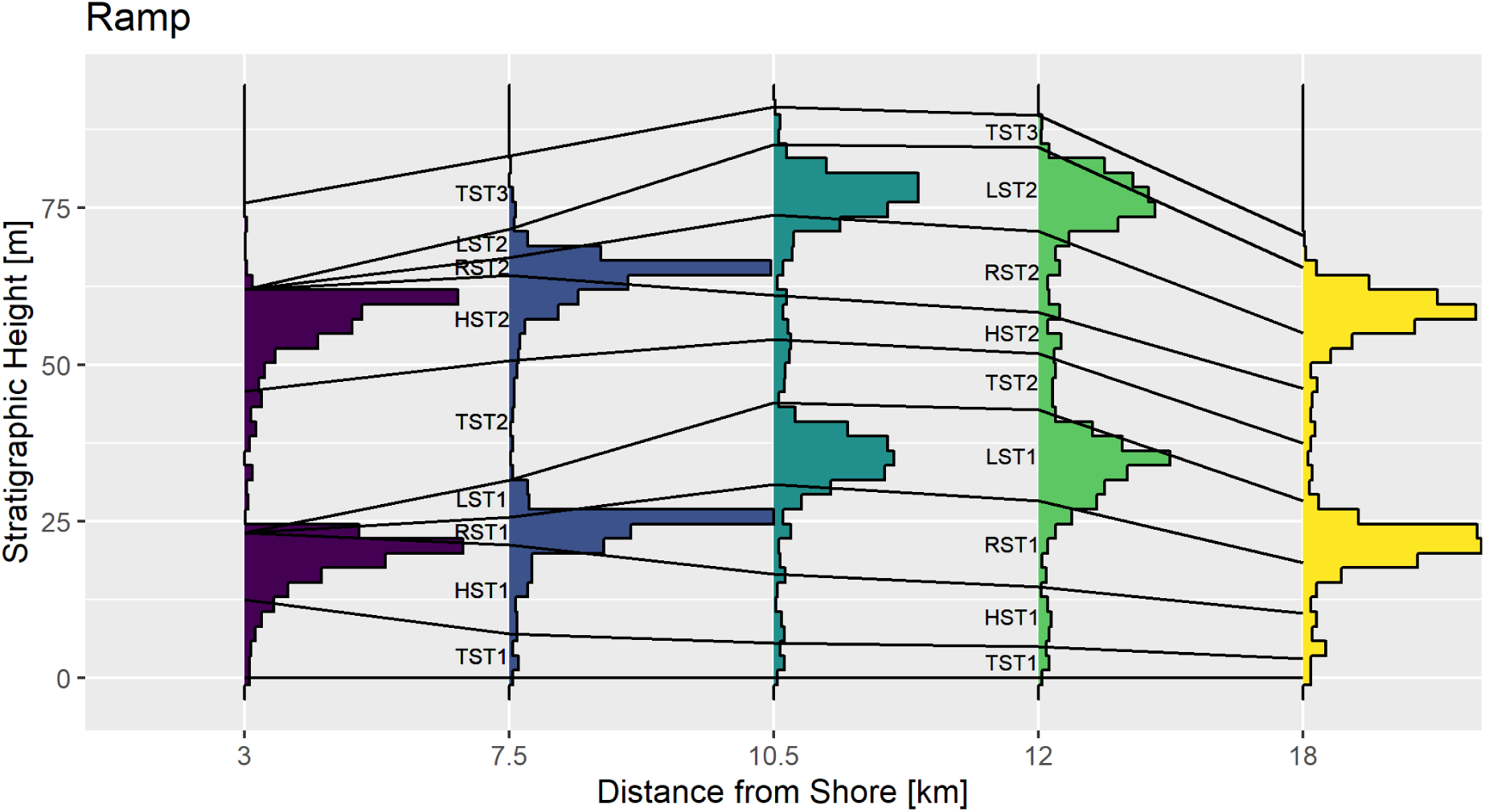
Spatial correlation of a pulse of extinction during the lowstand systems tract (LST) in the ramp geometry, assuming a sampling rate of 10 fossils per Myr. See Supplementary Figures 18 and 19 for spatial correlation of other extinction scenarios.

### Range offset

We used range offset to quantify the joint effects of ecology, abundance, and condensation and unconformity bias on biostratigraphic precision (Holland, 2000; Holland and Patzkowsky, 2002). We distinguish between two types of range offset:

1. Temporal range offset, defined as the age difference between the last occurrence of a taxon and its true extinction (measured in Myr),
2. Stratigraphic range offset, defined as the height difference between the location of the last occurrence of a taxon and the horizon of its true extinction (measured in m) (Figure 10, Table 4). For each geometry and examined position, we determined the (local) range offset of 1000 randomly generated taxa based on a constant extinction rate using the “range_offset” function from the StratPal package. Each taxon is determined by its fossil sampling rate and niche. Sampling rates were drawn from a uniform distribution on a log scale with values between 2 and 100 fossils sampled per Myr. Niches were specified via the probability of collection model by Holland and Patzkowsky (1999) as implemented by the function “snd_niche” in the StratPal package and using water depth as gradient. Preferred water depth was drawn from a uniform distribution with minimum and maximum values of 0 and 120 m, respectively. Water depth tolerance was drawn from a gamma distribution with shape parameter of 1 and scale parameter of 10. Accordingly, water depth tolerance has both a mean and standard deviation of 10 m.

**Figure 10:**
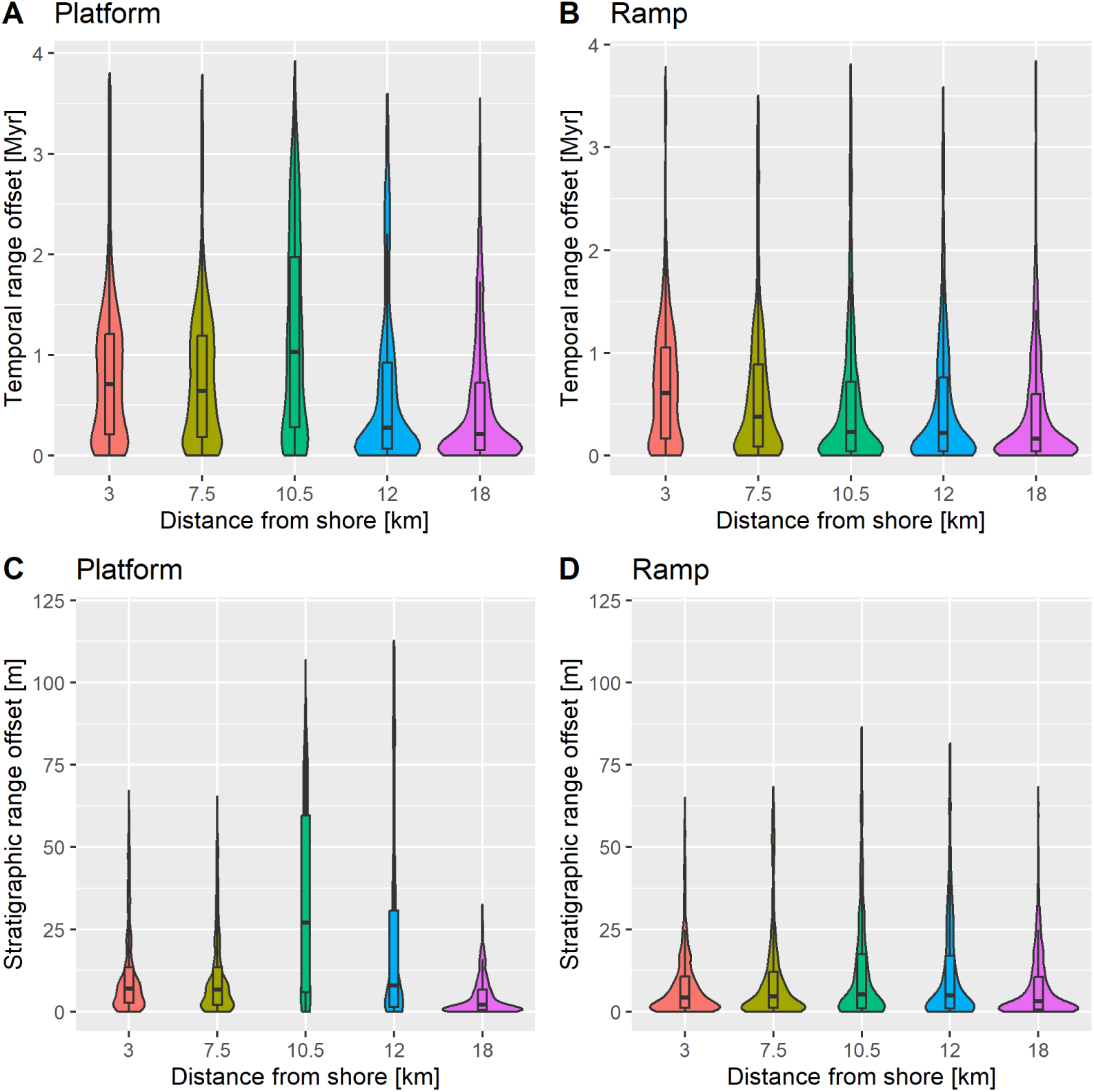
Temporal range offset (top row, A and B) and stratigraphic range offset (bottom row, C and D) in the platform and ramp geometries. Boxplot hinges are the 1^st^ and 3^rd^ quartile, center is the median. Summary statistics are given in Table 4.

**Table 4:** Median and 1^st^ (Q1) and 3^rd^ (Q3) quartiles of temporal and stratigraphic range offset in the platform and ramp geometries. Temporal range offset is rounded to 10 kyr, stratigraphic range offset to decimeters.

| Geometry |  | Platform |  |  |  |  | Ramp |  |  |  |  |
| --- | --- | --- | --- | --- | --- | --- | --- | --- | --- | --- | --- |
| Distance from shore [km] |  | 3 km | 7.5 km | 10.5 km | 12 km | 18 km | 3 km | 7.5 km | 10.5 km | 12 km | 18 km |
| Temp. Range offset [Myr] | Q1 | 0.21 | 0.18 | 0.28 | 0.07 | 0.05 | 0.16 | 0.09 | 0.04 | 0.04 | 0.04 |
|  | Median | 0.71 | 0.64 | 1.03 | 0.28 | 0.22 | 0.61 | 0.38 | 0.23 | 0.22 | 0.16 |
|  | Q3 | 1.21 | 1.19 | 1.97 | 0.92 | 0.72 | 1.05 | 0.89 | 0.72 | 0.76 | 0.6 |
| Strat. range | Q1 | 2.4 | 1.7 | 5.0 | 1.4 | 0.4 | 1.3 | 0.9 | 1.0 | 0.8 | 0.7 |
|  | Median | 6.5 | 5.7 | 25.4 | 7.5 | 1.8 | 4.2 | 3.7 | 4.7 | 4.2 | 3.5 |
|  | Q3 | 13.8 | 12.1 | 58.0 | 31.8 | 6.6 | 9.6 | 10.2 | 17.5 | 15.7 | 11.6 |

## Results

### Stratigraphic architectures

Although forced by identical sea level curves, the simulated stratigraphic architectures differ remarkably due to differences in sediment transport, carbonate factory production dynamics, and lithification time.

The platform geometry simulation results in a non-rimmed platform dominated by aggradation, as shown by the stacked block shape of the Wheeler diagram. The platform continuously progrades, with the platform edge moving 2 km slopewards over the course of the simulation (Figure 1, Supplementary Figure 3). In contrast, the ramp simulation produces a distally steepened ramp with progradation and retrogradation in phase with the 3^rd^ order sea level fluctuations, leading to a Wheeler diagram with lenticular bodies and no net progradation over the simulated 4 Myr (Figure 1, Supplementary Figure 4).

### Platform

Age-depth models on the distal and proximal platform top are indistinguishable and dominated by long gaps covering both the highstand and the regressive systems tracts (HST1, RST1, HST2 and RST2), with sediment being predominantly formed by the euphotic factory (Figure 1). Preservation of systems tracts differs across the prograding platform edge, with the proximal prograding platform edge preserving neither HST1, RST1, HST2, nor RST2, but the distal prograding platform edge fully preserving both HST2 and RST2. On the distal slope, the age-depth model is almost continuous, interrupted only by short gaps, with sedimentation rates fluctuating around 10 m/Myr. In the platform top, sediment accumulation peaks during the transgressive systems tracts. Generally, sediment accumulation averaged over 100 kyr is less than 20 % of the maximum growth rate of the dominant carbonate factory (Supplementary Figure 5). Stratigraphic completeness measured on the scale of 1 kyr (model resolution) is below 25 % on the platform top, increases to almost 100 % across the prograding edge, and drops to approx. 80 % on the distal slope (Supplementary Figure 6). Gap durations on the platform top reflect approximate half-periods of the 3^rd^ and 5^th^ order fluctuations in sea level, whereas gaps on the slope are dominated by short-term gaps at the scale of model resolution (Supplementary Figure 7).

### Ramp

Age-depth models on the inner ramp are similar to those on the platform top, with prolonged gaps associated with subaerial exposure and the resulting amalgamation of gaps produced by 3^rd^ and 5^th^ order sea level changes that fully remove HST1, RST1, LST1 and RST2. Distally, the influence of subaerial exposure weakens, with only parts of HST1, RST1, LST1 and RST2 missing in the middle ramp, and all systems tracts being preserved on the ramp slope and outer ramp. Generally, completeness on the timescale of 1 kyr is higher than in the platform simulation, with values of 40 % on the inner ramp monotonously increasing to the distal outer ramp, where they reach almost 100% due to continuous delivery of sediment from the inner ramp or the aphotic factory (Supplementary Figure 6). Generally, the ramp geometry is dominated by shorter gaps to a larger extent and only reflects 3^rd^ order sea level changes on the inner ramp (Supplementary Figure 7). The number of hiatuses is lower than on the platform, with a peak on the proximal and distal ramp slope (6 to 12 km), and is dominated by shorter intermittent transport events (Supplementary Figure 8).

Comparing the two simulations, the platform shows a clear separation into top and slope, with the prograding platform edge forming a narrow boundary between them. In contrast, the transition from onshore to offshore deposition on the ramp system is gradual, with age-depth models and water depths gradually changing along the onshore-offshore gradient (Figure 3). Proximally, section thickness in both simulations is subsidence-limited, i.e. denudation removes all subaerially exposed deposits, and record formation is thus limited by accommodation space. Thick sections are only formed on the prograding rim (prograding platform edge and ramp slope) where slope deposits accumulate fast enough to build up to water depths where euphotic and oligophotic carbonate producers pick up production and then rapidly fill available accommodation space (Supplementary Figure 9, Figure 3). In the platform, the majority of hiatuses are generally short (< 100 kyr) but with a long tail formed by hiatuses with durations longer than 1 Myr. These very long hiatuses are only present on the platform top and the prograding rim. In contrast, the median hiatus duration on the inner ramp is longer than 500 kyr, and then gradually tapers off towards the ramp slope, with very long hiatuses only present in the most proximal parts (Supplementary Figure 10). Generally, time is much more continuously distributed in the ramp compared with the platform (Supplementary Figures 5 and 11).

Denudation removes significant portions of sediment accumulated on the platform top, leading to the amalgamation of adjacent gaps and, as a result, removal of large parts of the highstand and regressive systems tract. This effect is weaker on the ramp due to lower precipitation (Figure 2 Table 2).

In summary, differences between simulations can be explained by differences in carbonate factory production, carbonate factory properties, sediment transport, and cementation. In the platform simulation, the highly productive euphotic carbonate factory, rapid cementation, and reduced transport leads to a clear “Winner takes it all” dynamics of platform growth (Schlager, 2005) that separates platform top from slope. The progradation speed of the platform rim is limited by slope deposits building up to water depths where the euphotic carbonate factory takes over carbonate production. Sea level drops lead to instantaneous subaerial exposure of the flat platform top and the formation of associated long hiatuses. This effect is reinforced by denudation that merges gaps resulting from successive sea level drops. On the slope, sedimentary dynamics are dominated by short, intermittent transport events and sedimentation from the aphotic factory. In the ramp simulation, the oligophotic carbonate factory has been set to be the most productive and accumulates sediment at greater depths than it is the case on the platform. Together with increased sediment transport and reduced cementation, this facilitates lateral distribution of sediment and suppresses the formation of a clear platform-like geometry, especially as the sea level amplitude is not large enough to generate water depth differences where oligophotic production drops significantly. Combined, this leads to a less pronounced spatial separation along the onshore – offshore gradient and shorter gaps.

### Ecological bias

Water depth (Figure 3, Supplementary Figure 12) on the platform top is shallow throughout as the euphotic carbonate factory quickly fills all available accommodation space. Across the prograding edge there is a shallowing upward trend with the deposits migrating from slope to platform top, while the distal slope deposits slowly drown (Figure 3 A). In the ramp simulation, differences in water depth across the sampling locations are more gradual, with the very shallow deposits of the platform top missing (Figure 3 B, Supplementary Figure 11). Generally, we observe no rapid changes in water depth across unconformities due to high rates of in situ sediment production that quickly fills all available accommodation space after re-flooding.

### Unconformity and condensation bias

Condensation ratio (Figure 4) is close to one throughout sections on the distal slope of the platform and the ramp slope and outer ramp, indicating that these locations are only weakly influenced by condensation and unconformities. In contrast, condensation ratios range from below 0.3 to 10 on the prograding platform edge, showing that constant rates and fluxes in the time domain will fluctuate by almost two orders of magnitude in the stratigraphic domain. As with the age-depth models, the platform displays a clear separation into high age misassignment bias on the top and no age misassignment bias on the slope, while age misassignment bias gradually decreases distally in the ramp.

### Preservation of extinction pulses

While extinction pulses can recognized and correlated laterally, they are preferentially recovered from the transgressive systems tract (TST) with a temporal misassignment of more than 1.5 Myr (two-thirds of the period of the 3^rd^ order sea level change).

### Abundance effects

At the prograding platform edge, the backwards-smearing, or the Signor-Lipps effect, is the most pronounced for intermediate sampling rates (10 and 22 fossils per Myr, Figure 6). When fossils are very abundant, last occurrences cluster at the unconformities been RST1 and LST1, as well as between TST2 and LST2, thus emphasizing the stratigraphic bias. In contrast, when fossils are rare, none of the long gaps are reflected (Figure 6). In the absence of stratigraphic biases (e.g., on the distal slope), backwards smearing is absent independently of sampling rate (Supplementary Figures 14 and 15). This clearly demonstrates that abundance bias is a secondary effect that compounds on other biases and can modify them in either direction.

### Extinctions by systems tracts

Comparing preservation of extinction pulses associated with specific systems tracts and a baseline scenario with constant extinction rate, we observe substantial backwards-shifting of last occurrences into stratigraphically lower systems tracts, most significantly the transgressive systems tract (Figure 7). For example, on the platform top, extinctions during the HST, RST, and LST will all be recovered from the TST on the platform top, thus clearly favoring the recovery of mass extinctions from records formed during sea-level rise. For extinction pulses associated with the TST we find the opposite effect, whereby the rapid accumulation of sediment during the TST can effectively dilute the extinction signal and make the extinction appear smoothed out, resulting in underestimation of extinction rates. Generally, preservation of extinction pulses is better in distal parts of the platform and ramp (Supplementary Figures 16 and 17).

### Spatial correlation of extinction pulses

Extinction pulses can generally be traced laterally throughout the simulation even if they are preserved in different systems tracts (Figure 8, Figure 9, Supplementary Figures 18 and 19). As a result of unconformity and condensation biases, extinction pulses show a slow onset and rapid decrease proximally, whereas the symmetric increase and decrease in extinction rates is well-preserved distally (Figure 9, Supplementary Figure 13). In the ramp, correlation is facilitated by the gradual transition between onshore and offshore deposits, where the RST and LST thin out proximally (Figure 9). In contrast, correlation between platform top and distal slope in the platform geometry is challenging due to the heterogeneous preservation of systems tracts across the prograding platform edge. For example, the proximal prograding platform edge preserves HST1 and RST1, but not HST2 and RST2. Across both geometries, extinction pulses during the TST are the easiest to trace laterally due to rapid sediment accumulation in this systems tract. In contrast, extinctions during the RST and LST move into the TST (platform) and HST (ramp) as sections become more proximal (Supplementary Figure 18, Supplementary Figure 19). Extinctions during the HST are recorded in the correct systems tract in the ramp, but will be recovered from the TST on the platform top due to removal of previously deposited sediment (Figure 2).

### Range offset

Biostratigraphic precision as measured by median temporal range offset on the platform top is approximately 700 kyr, peaks at the proximal prograding platform edge with a median of 1.04 Myr (3^rd^ quartile: 1.97 Myr) between the true disappearance of a taxon and its last occurrence, and drops almost fivefold on the slope with median values of 280 and 220 kyr (Figure 10 A, Table 4). Reduced biostratigraphic precision across the prograding platform edge is most likely due to ecological bias resulting from the shallowing-upward trend due to the progradation of the platform. Range offset on the platform top is driven by stratigraphic and abundance bias, as water depth is shallow throughout the simulation (Figure 3 A). On the ramp, temporal range offset decreases gradually across the onshore-offshore gradient, with median values ranging from 600 kyr (inner ramp) to 160 kyr (distal outer ramp). This is a result of decreasing gap duration that is not fully counteracted by a moderate increase in water depth fluctuations (Figure 3).

Stratigraphic range offset in the ramp is moderate, with a median of around 4 meters between the last occurrence of a taxon and its true extinction horizon (interquartile range approx. 10 m). Interquartile ranges are slightly elevated on the proximal and distal ramp slope, most likely due the increased thickness accumulated in the prograding/retrograding band (Supplementary Figure 9) in combination with increases in temporal range offset. Similarly, stratigraphic range offset in the platform geometry peaks on the proximal prograding platform edge (median 25 m) and is elevated on the distal prograding platform edge (median 7.5 m, 3^rd^ quartile 31.8 m) (Figure 10, Table 4).

## Discussion

Comparing the stratigraphic distribution of fossils between carbonate platforms and ramps, we found clear partitioning of the carbonate platform into shallow-water platform top deposits dominated by long hiatuses and quasi-continuous slope deposits with larger variations in water depth. Carbonate ramp deposits display a gradual transition along the onshore-offshore gradient, with distally increasing variability in water depth and diminishing influence of hiatuses. Platform top and inner ramp deposits will thus be dominated by unconformity bias, whereas ecological biases due to niche tracking become increasingly important on platform slope and distal ramp deposits (Figure 1). These contrasts between geometries are determined by differences in carbonate factory productivity and production profile, as well as by sediment transport and cementation. Our results indicate that understanding the response of carbonate factories to external forcing predicts the structure of marine carbonate stratigraphic architectures and can thus provide useful first-order approximations on their stratigraphic biases and how these propagate into downstream study systems that rely of fossil data (e.g., extinction and origination dynamics, climate change, and phenotypic evolution).

### Carbonate ramps vs platforms

Here we have focused on the differences between a tropical carbonate platform and a temperate ramp, thus highlighting how depositional systems with identical external forcing record earth system signals depending on their latitude and their constituting carbonate factories. Instead of reproducing specific existing carbonate platforms, the simulated scenarios represent conceptual endmembers of carbonate geometries. Latitude is not the sole determinant of carbonate depositional geometry: Pomar (2001) reported the transition from a ramp to platform geometry due to an ecological succession from grain- to framework producing biota, while Lehrmann et al. (2022) attributed a transition from ramp to platform to cement stabilization of margin deposits due to increased carbonate saturation of sea water. In contrast, transitions from platforms to ramps are commonly associated with major perturbations of the biogeosphere, e.g., ramps are more abundant than platforms following major mass extinctions (Burchette and Wright, 1992). Kammer and Ausich (2006) explicitly link changes is crinoid diversity to the collapse of coral-stromatoporoid carbonate platforms into ramps during the Late Devonian mass extinction, and there is ample evidence for a decrease in framework building organisms and dominance of microbial carbonate production after mass extinctions (Mata and Bottjer, 2012; Ibarra *et al*., 2016; Yao *et al*., 2016). Over macroevolutionary timescales, varying groups of carbonate producing organisms become dominant, potentially leading to the co-evolution of life and its record. However, these effects are beyond the lifespan of an individual carbonate ramp, and Pomar (2001) argues that, although the organisms in carbonate depositional ramps varied drastically throughout the Earth’s history, they play similar roles in the formation of carbonate ramps. Using exploratory simulations, we found that transition from platform to ramp geometry is initialized by backstepping on the platform top and increased deposition on the distal slop to achieve the more gradual ramp profile (Supplementary Figure 20), while transitions from ramp to platform geometry result in rapid progradation as the euphotic factory rapidly picks up production on the ramp slope (Supplementary Figure 21). This indicates that fossil records of perturbations of the biogeosphere will have a distinct stratigraphic overprint resulting from transitions between carbonate platform geometries.

Transitions between carbonate geometries highlight that carbonate platforms are hysteretic systems whose architecture depends not only on external forcing but also on their accumulative history and internal feedbacks. In our results, this is exemplified by the prograding platform edge, where, depending on the advancement of the progradation, the HST and RST are preserved or not and extinction pulses will accordingly be recovered from different systems tracts (Figure 8).

Despite high carbonate growth rates resulting in keep-up sedimentation that closely tracks sea level, drowned carbonate platforms are common in the geological record. Potential explanations for this apparent paradox include drastic reduction in benthic production or rapid generation of accommodation space (e.g., due to sea level rise or tectonics) that outpaces benthic carbonate production (Schlager, 1981; Szulczewski, Belka and Skompski, 1996; Petrovic *et al*., 2023). Drowning behavior depends on the relative importance of transport vs. in-situ production and can result in back-stepping or pinnacle geometries, respectively (Wang, Burgess and Rankey, 2026). As a first-order approximation, our results suggest that the deepening-upward trend within a drowning platform will result in decreasing influence of unconformity bias, increasing the contribution of condensation bias in parallel to carbonate production decrease and an ecological transition from shallow- to deep-water taxa.

### Ecological bias and water depth as principal ecological gradient

In alignment with the majority of the literature on marine stratigraphic paleobiology (Holland 2000; Nawrot et al. 2018), we have adopted water depth as the principal ecological gradient in shallow-marine environments. This is based on the argument that (1) water depth correlates well with complex biologically important parameters such as oxygen and light availability, and (2) variations in water depth will translate into empirically recognizable variations in lithofacies. We found little to no variations in water depth on the platform top and the proximal ramp, as carbonate production closely tracks the sea level as accommodation space becomes available. The prograding platform edge shows a shallowing-upward trend, resulting in stratigraphically increasing importance of unconformity biases, whereas the distal slope slowly drowns (Figure 3, Supplementary Figure 12). Distally, water depth is in 1-1 correspondence with the eustatic sea level curve, in a striking contrast with results from siliciclastic systems (see e.g. Figure 5 in Holland and Patzkowsky (2015)). We observed no rapid change in water depth across unconformities (Figure 3), suggesting that ecological and unconformity bias do not compound in carbonate systems as they do in siliciclastic systems (Nawrot et al. 2018; Holland and Patzkowsky 2015). These differences are rooted in the high rates of in-situ keep-up sedimentation in carbonates, as opposed to the point-source sediment distribution in siliciclastics. Ecological bias introduced by changes in water depth is only recognizable on the prograding platform edge (10.5 and 12 km from shore), where extirpation of deeper-water taxa due to a shallowing-upward trend inflates range offset (Figure 10, Table 4).

Empirically, in shallow water (< 40 m) carbonate systems over short spatial scales (< 10 km), water depth ceases to become a reliable predictor of facies type (Ginsburg, 1956; Rankey, 2004; Purkis, Rowlands and Kerr, 2015). Additionally, key environmental parameters such as salinity and hydrodynamic energy can vary drastically shallow water depositional environments such as lagoons. For example, recent work in the Bahamas suggests that the distribution of organism habitats are influenced by complex interactions of water-depth, hydrodynamic-settings, incumbency, and seabed sediment type (Liu *et al*., 2026). Together, this suggests that water depth ceases to be a useful principal ecological gradient in shallow-water carbonate systems, and should instead be replaced by either complex compound gradients or facies categories identifiable in the rock record. The combination of spatially complex facies mosaics and rapid in-situ keep-up sedimentation in tropical carbonate deposits suggests that results on niche tracking from siliciclastic deposits might not easily generalize to carbonates (Brett *et al*., 2007).

### Unconformity and condensation biases and the common cause hypothesis

By quantifying unconformity and condensation bias with the condensation ratio (equation 1), we found that rates observed in the stratigraphic domain can differ by almost two orders of magnitude relative to the true rates in the time domain when the underlying true age-depth relationships are not correctly resolved (Figure 4). This effect only depends on the misassignment of ages to stratigraphic positions (Figure 2) and will accordingly propagate downstream into all rates observable in the stratigraphic domain that are reconstructed from fossils (e.g., extinction and origination rates and rates of phenotypic evolution) (Hohmann, 2021). While rates are overestimated 10-fold across the unconformity separating the TST and the LST and underestimated more than 3-fold within the rapidly accumulating TST on the platform top (Figure 4 A), rates on the distal outer ramp can be recovered almost without stratigraphic inflation or deflation as the condensation ratio remains approximately 1 throughout the section (Figure 4 B). As observed by Hohmann et al. (2024), this favors the recovery of complex evolutionary dynamics from the platform top and proximal ramp when the underlying mode of evolution in the time domain is gradual.

Red flags for stratigraphically inflated rates are (1) elevated rates associated with indicators of sea level drops (e.g., subaerial exposure or facies shifts indicating a deepening), (2) simultaneous elevation of potentially independent rates, and (3) distally decreasing rates when correlating through the basin. However, these red flags may be genuine hypotheses; for the examples above, one could consider that extinction rates are higher during eustatic sea-level falls owing to the loss of habitat; potentially independent rates varying in concert might be taken to indicate an unknown ecological dependency, and evolutionary rates might truly vary along a water depth gradient. Furthermore, environmental factors – including sea level – are commonly reconstructed from fossil data. These dependencies between the physical and fossil records pose a risk of circular reasoning, whereby the biological rate shifts and the environmental change are used to mutually interpret each other (Brett, 1998). Stratigraphic paleobiology largely focuses on how stratigraphy overprints (presumably independent) paleontological signals. The common cause hypothesis extends this to scenarios where the linkage of earth systems drives both paleontological and stratigraphic patterns (Peters and Heim, 2011). Polly (2025) provided an illustration of the common-cause hypothesis by demonstrating that spatially heterogeneous sedimentary systems can generate complex evolutionary patterns mimicking punctuated equilibrium even in the absence of true speciation. This demonstrates that the causal relationships between external forcing and internal feedbacks of sedimentary systems and their records can be complex and multidirectional, and might not be testable with single-section observations. Furthermore, this study emphasized the importance of spatial heterogeneity in driving and identifying environmental and evolutionary shifts – a feature particularly prominent in carbonate depositional systems (Liu et al. 2026). The fossil record offers the opportunity to separate common cause from correlation or internal feedback. The integration of stratigraphic forward models with biological simulations allows us to formulate multiple alternative scenarios, which can guide spatial sampling of empirical data and which can then be assessed in terms of how well they explain that data.

### Preservation of extinction pulses and abundance bias

Four of the big five mass extinctions are associated with rapid changes in sea level, resulting in patterns of last occurrences that are a mixed signal of stratigraphic and biotic effects (Holland and Patzkowsky 2015; Holland 2020). Using simulations of siliciclastic systems, Holland and Patzkowsky (2015) argue that clustering can be attributed to two factors: Subaerial unconformities due to drops in sea level and stratigraphic condensation across major flooding surfaces due to abrupt deepening. Our results show clustering of last occurrences in the transgressive systems tract (rising sea level) on the platform top below the subaerial unconformity when the extinction rate is constant or when its pulses are associated with the HST, RST, or LST (Figure 7). However, we find no indication of stratigraphic condensation as expressed by depressed sedimentation rates (Supplementary Figure 5). We attribute this to high rates of in situ carbonate production resulting in predominantly aggradational sediment accumulation, contrasting the sediment starvation associated with the rise of the erosional base during TST in siliciclastic systems. Zimmt et al. (2021) showed that stratigraphic biases can be mitigated by basin-wide analyses of last occurrences and coarsening of the stratigraphic resolution. Extinction pulses can generally be correlated laterally in our results (Figure 8, Figure 9), though they will preferentially be recovered from the TST, with the RST and HST fully missing from platform top deposits. While spatial correlation of extinction pulses might be effective in ramp geometries where systems tracts gradually pinch out proximally, this approach might not be feasible in platform geometries, where extinction pulses are offset by up to 1.5 Myr and three systems tracts (Figure 7). Holland (2020) argues that stratigraphic biases substantially shorten the perceived duration of mass extinctions. We find the opposite for extinctions associated with rises in sea level, where extinction rates are effectively diluted threefold due to rapid carbonate platform growth during the TST (Figure 4, Figure 7).

We found that the Signor-Lipps effect (i.e., the backwards smearing of extinction pulses due to low fossil sampling rates (Signor *et al*., 1982)) carries the highest risk of wrong identification of extinction pulses when sampling rates are intermediate (10 to 22 fossils per Myr): If fossils are more abundant, clusters of last occurrences are closely associated with hiatal surfaces, making it easy to recognize them as stratigraphic artefacts. In contrast, if fossils are very rare, backwards smearing is strong enough to remove the stratigraphic bias in total. For intermediate sampling rates, artefactual peaks in last occurrences can appear detached from their causal hiatal surface only when fossils are rare enough to lead to a significant backwards smearing from the hiatal surface but abundant enough to preserve the peak in last occurrences (Figure 6). Abundance bias introduced by varying fossil sampling rates is thus context-dependent and can both pronounce and diminish other biases. For comparison with previous studies in stratigraphic paleobiology (Holland, 1995; Holland and Patzkowsky, 2015), we have focused on the Signor-Lipps effect sensu stricto. However, our results will hold mutatis mutandis to the forwards smearing of origination rates (Foote, 2001).

Generally, fossil abundance has stratigraphically downstream effects on all statistical analyses by reducing sample size and, as a result, statistical power. Abundance bias thus constitutes a second order effect whose impact depends on the research question. For example, Hannisdal (2006) found that small sample size due to non-preservation leads to the erroneous rejection of null hypotheses and identification of stratophenetic stasis. In empirical studies, the number of fossil specimens is known at the analysis stage. Combining synthetic fossil records simulated under specific hypotheses and with the available sample size allows to establish a baseline for best-case performance of statistical methods (Hohmann *et al*., 2024). Because of this analytical traceability, we believe that abundance bias is the least concerning of the biases discussed here.

### Range offset

We found that biostratigraphic precision as measured by temporal range offset is commonly below the precision of geochronological approaches, with median values of up to 1 Myr on the distal prograding platform edge (Figure 10, Table 4). Although our values of range offset are not directly comparable to those in Holland and Patzkowsky (2002) due to differences in simulation setup and parameters, we arrive at comparable orders of magnitude. Empirically, range offset of the order of 100s of kyrs is common even for abundant microfossils in deep-sea environments (Dowsett, 1988; Miller *et al*., 1994; Kucera and Kennett, 2000), supporting that our values of range offset are plausible approximations for the biostratigraphic precision achievable in carbonate platforms. Range offset depends on fossil abundance, unconformity, condensation, and ecological biases, making it a priori the most complex statistic examined in this study. However, the structure of carbonate platforms provide heuristics on range offset. For example, on the platform top, water is shallow throughout and long gaps are present, and range offset will accordingly be controlled by unconformity and abundance bias. On distal parts of the slope, range offset will be dominated by fossil abundance as gaps are short and rare with moderate fluctuations in water depth. Range offset is highest in prograding parts of the platform where unconformity bias and ecological bias compound due to the joint influence of long gaps and shallowing-upward trends. Answering palaeobiological questions using fossils from strata with irregular age-depth relationships requires precise, unbiased estimates of said age-depth relationships. Our results indicate that in such strata, age-depth models constructed from biostratigraphy will be most biased, even for abundant eurytopic taxa. A way to counteract this would be to incorporate methods that estimate range offset (e.g., Schueth et al. (2014)) or forward models as inference tools into the construction of age-depth models.

Stratigraphic range offset (distance between the last occurrence and the true extinction horizon) is multiple meters in our simulation setup, with a peak at the proximal prograding platform edge where it reaches median values of 25.4 m. While this implicitly follows from the results on temporal range offset, it highlights that last occurrences can be separated from their associated extinction horizons by tens of meters, especially in the rapidly accumulating TST.

### Model limitations

The used simulation setup is sequential, with the palaeobiological simulation following the stratigraphic simulation without interacting with it. Specifically, while CarboKitten has a complex sediment transport model based on the active layer approach proposed by Paola et al. (1992), fossils in our setup are not transported laterally or reworked by entrainment, erosion, or bioturbation.

Empirically, most fossil assemblages are representatives of their local communities and transported only short distances, with exceptions being identifiable by their taphonomic condition (Kidwell and Flessa, 1995), and Patzkowsky and Holland (2012) argue that lateral transport is negligible relative to other stratigraphic biases. Tighter integration of biological and sedimentological simulations would allow to explore this effect in silico or to calibrate the transport model.

The effective depositional resolution in our simulations is 1 kyr (model resolution), meaning we assume records are continuous on a millennial timescale due to time-averaging of sedimentary particles even if sediment deposition is intermittent (Kowalewski and Bambach, 2008). Time-averaging in many depositional systems is multiple kyr, though much lower values have been observed when sedimentation is rapid (Tomašových *et al*., 2018). Additionally, differences between carbonate and siliciclastic systems are reported, potentially due to differences in cementation, dissolution, and particle transport (Kidwell, Best and Kaufman, 2005; Kosnik *et al*., 2009; Kowalewski *et al*., 2017; Slootman *et al*., 2026b, 2026a). While we use internal time-steps of 100 years for our CarboKitten simulation (with potentially even smaller increments used to solve the underlying differential equations), reporting at this precision is not appropriate given the limitations of temporal resolution introduced by the physical process of sediment deposition and mixing.

## Conclusions

Combining forward simulations of attached carbonate platforms with spatially explicit models of fossil occurrences, we found that platform geometries show clear separation into platform top deposits with strong unconformity bias and quasi-continuous slope deposits. In contrast, carbonate ramp successions show gradually diminishing unconformity bias along the onshore-offshore gradient. The influence of ecological bias is weak due to high rates of in-situ keep-up sedimentation. These biases will propagate into downstream analyses and affect all study systems that draw information from the fossil record (e.g., paleoecology, evolutionary biology, and paleoclimatology). Extinction pulses associated with different systems tracts are easier to correlate on carbonate ramps and preserved in downdip locations, allowing to distinguish artefactual peaks in last occurrences introduced by unconformities from true extinction pulses. Differences between carbonate platform architectures can be attributed to variations in carbonate factory composition and productivity, sediment transport dynamics, and rates of cementation. Large-scale perturbations of the biogeosphere can modify the composition of carbonate factories and, by extension, stratigraphic architectures and the signature of the perturbation itself.

Our results show that earth system records differ predictably within and between depositional systems. Comparison of earth system records across depositional systems and combination of empirical studies with forward simulations as qualitative or semi-quantitative baselines are promising venues to separate earth system signals from overprints introduced by stratigraphy and bridge the simulation-empiricism gap in stratigraphic paleobiology.

## Code and data availability

All code and data is available under https://doi.org/10.5281/zenodo.20670764 (Hohmann, Jansen, *et al*., 2026), supplementary materials are available under https://doi.org/10.5281/zenodo.20671213 (Hohmann, Bickerton, *et al*., 2026).

## Acknowledgements

We would like to thank Johan Hidding (Netherlands eScience Center) for support with CarboKitten functionality and Ton Markus (Utrecht University) for help with preparing figures.

## Author contributions

NH: Conceptualization, Formal Analysis, Investigation, Methodology, Software, Supervision, Validation, Visualization, Writing (original draft, review and editing). SB: Investigation, Methodology. AJ: Investigation, Methodology. XL: Writing (review and editing), Methodology. EJ: Conceptualization, Funding acquisition, Validation, Supervision, Writing (review and editing).

